# Islet primary cilia motility controls insulin secretion

**DOI:** 10.1101/2021.12.14.472629

**Authors:** Jung Hoon Cho, Zipeng A. Li, Lifei Zhu, Brian D. Muegge, Henry F. Roseman, Toby Utterback, Louis G. Woodhams, Philip V. Bayly, Jing W. Hughes

## Abstract

Primary cilia are specialized cell-surface organelles that mediate sensory perception and, in contrast to motile cilia and flagella, are thought to lack motility function. Here we show that primary cilia in pancreatic beta cells exhibit movement that is required for glucose-dependent insulin secretion. Beta cell cilia contain motor proteins conserved from those found in classic motile cilia, and their 3D motion is dynein-driven and dependent on ATP and glucose metabolism. Inhibition of cilia motion blocks beta cell calcium influx and insulin secretion. Beta cells from humans with type 2 diabetes have altered expression of cilia motility genes. Our findings redefine primary cilia as dynamic structures possessing both sensory and motile function and establish that pancreatic beta cell cilia movement plays a critical role in controlling insulin secretion.

## MAIN TEXT

Cilia are slender, hair-like projections that mediate cellular interactions with the surrounding environment. There are two main classes of vertebrate cilia: primary cilia, which are solitary, present on most cells of the body, and specialize in sensory and signaling function (*1*), and secondary (motile) cilia which are present in large numbers per cell in select tissues and beat in coordinated waves (*2*). Fundamentally, these two cilia types are distinguished by their capacity for movement. Classic motile cilia possess a “9+2” axonemal microtubule structure, composed of a ring of nine outer microtubule doublets and a central pair, with interconnecting motor protein complexes such as dynein arms and radial spokes (*2, 3*). In contrast, conventional primary cilia possess a “9+0” configuration of microtubules (*4, 5*) and are considered immotile due to the lack of central pair and motor accessories.

There are exceptions to this dichotomous cilia classification, most notably the embryonic nodal cilia, which are “9+0” primary cilia that possess both motility and signaling capacity (*6*–*8*). In other tissues, primary cilia can bend passively under external forces such as fluid flow (*9*), and cilia mechanosensing regulates cellular homeostasis via ciliary and cytoplasmic Ca^2+^ (*10*–*12*). Primary cilia can also exhibit spontaneous fluctuations from internal actin-myosin forces generated at the cell cortex (*13*), but active dynein-driven motility in primary cilia has not been reported. Ultrastructural studies have shown that primary cilia architecture can differ extensively from the classic “9+0” paradigm, including functional central pairs (“7+2”), microtubule rearrangements, and protein densities along the axoneme mimicking motile structures (*14*–*20*). Collectively, these emerging data suggest that primary cilia may in fact be dynamic and that the distinction between primary and motile cilia may not be absolute.

To investigate the possibility of primary cilia being motile and the functional significance of their motility, we used live imaging to examine primary cilia behavior *in situ* in pancreatic islets. Defects in primary cilia cause metabolic disorders seen in human ciliopathy syndromes (*21*), and polymorphisms in cilia genes are linked to human obesity and type 2 diabetes (*22, 23*). Pancreatic beta cells secrete insulin in response to extracellular glucose, a process regulated by primary cilia and ciliary GPCRs (*24*–*27*). But what allows beta cells to respond to glucose quickly and *en masse*, despite their heterogeneity and fixed distribution within the islet? We hypothesized that, in the relatively enclosed islet environment where beta cells themselves are nonmotile, cilia projecting from their surface can nonetheless move to seek ambient signals, and that this motility would contribute to an orchestrated and more rapid whole-islet response. Here we show that beta cell cilia indeed possess active motility, driven by dynein and ATP, which is required for coordinated calcium influx and insulin release. These findings identify cilia motility as a new core beta cell function that underlies hormone secretion.

## RESULTS

To study real-time behavior of primary cilia in islet beta cells, we generated beta cell cilia GFP reporter mice by crossing INS-1 Cre (*28*) with ciliary somatostatin receptor 3 (SSTR3) GFP-OFF mice (*29*) (**Figure 1A**). The INS1-Cre strain was chosen for its efficient and selective beta cell recombination without ectopic expression in the central nervous system (*28*). Homozygous beta cell cilia GFP mice are fertile and produce normal size healthy litters, expressing constitutive green fluorescence in the primary cilia of beta cells. Fluorescence is robust and withstands prolonged live imaging and fixation (**Figure 1B**). Ultrastructural examination of the beta cell cilium reveals a rod-like axoneme emanating from near the nucleus, composed of microtubule filaments arranged cylindrically and symmetrically with less-defined electron densities connecting the microtubules. A pair of orthogonally arranged centrioles are found in a neighboring cell, composed of longitudinal filaments, amorphous luminal densities, and external appendages (**Figure 1C**).

**Figure 1.**
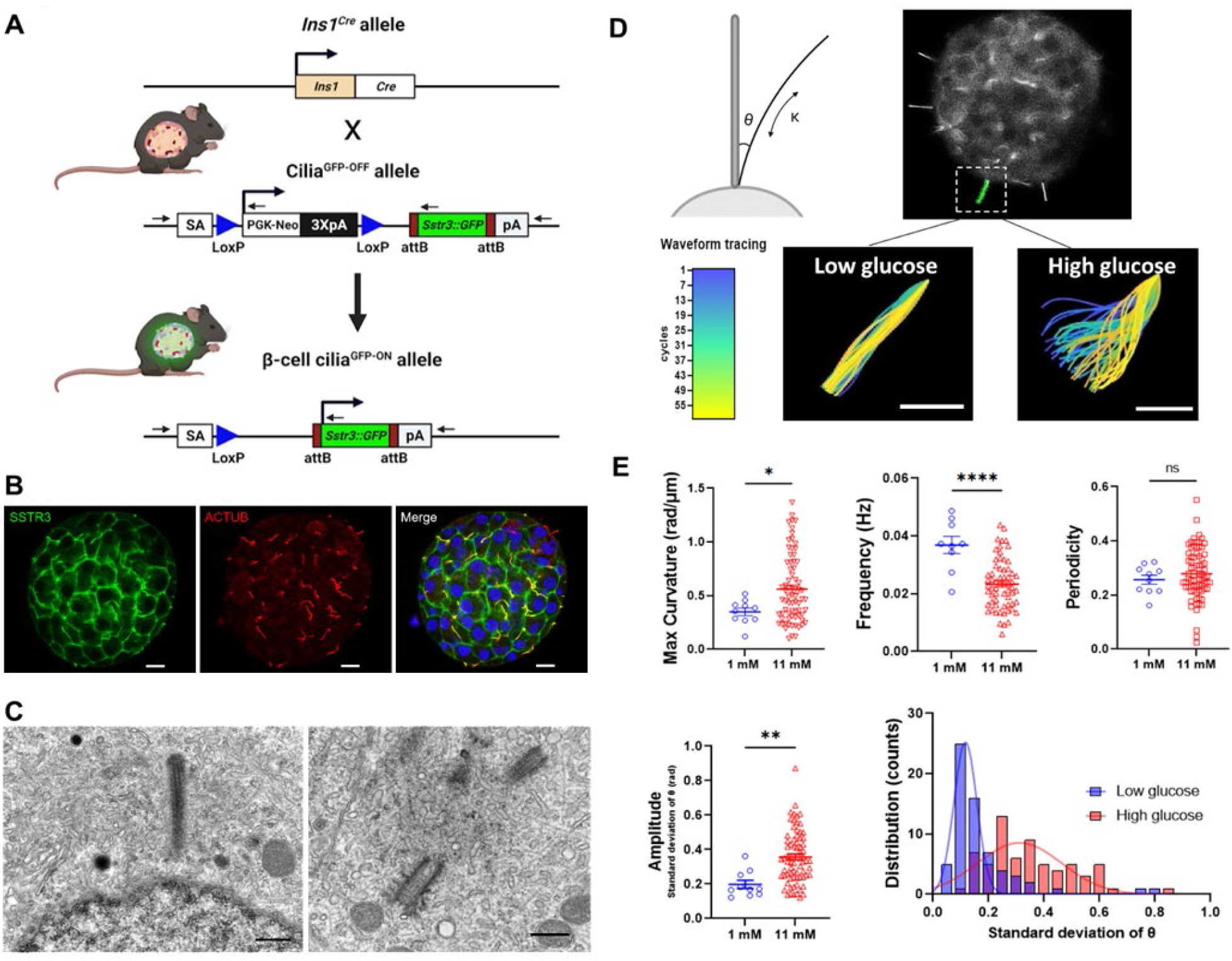
Beta cell cilia exhibit glucose-dependent motility. (**A**) Beta cell cilia reporter mice were generated by crossing INS1-Cre with SSTR3-GFP^OFF^ strains. (**B**) SSTR3 protein is enriched in mouse beta cell cilia with a small amount of detectable expression in the plasma membrane. Co-localization of SSTR3 (green) with acetylated alpha tubulin (cilia marker, red) is seen as yellow in the merged image. Non-beta cell cilia are red-only. Nuclei are labeled in DAPI, blue. Scale, 10 µm. (**C**) Transmission electron micrograph of human beta cell cilium (left, scale 500 nm) and a pair of centrioles (right, scale 400 nm). (**D**) Beta cell cilia exhibit glucose-dependent motility. Schematic shows measurement of theta (θ) as the angle of deflection and kappa (κ) as the curvature of the cilia axoneme. Video microscopy captures primary cilia movement on murine beta cells in response to glucose (**Movie S1**), with representative traces shown in colored panels corresponding to the temporal heatmap where the starting frame of the ciliary wave is depicted in blue and the end in yellow. Scale, 5 µm. (**E**) Cilia waveform analysis showing increased ciliary curvature, reduced frequency, and increased amplitude at high glucose (**Movie S1**). *p < 0.05, * *p < 0.01, * * *p < 0.001. Histogram shows cilia amplitude distribution at low (blue) versus high (red) glucose.

Live cilia imaging revealed spontaneous movement of beta cell cilia in intact islets. Cilia at low glucose displayed slow and variable oscillations, with wave periods on the order of 20-30 seconds (**Movies S1**). This contrasts with the fast and regular beating of conventional motile cilia with periods in the 0.1-0.3 second range (*30*–*32*). In addition to planar beating, beta cell cilia displayed a lasso-like 3D rotation, whose trajectory was less amenable to tracing as it moved in and out of imaging focus and required a z-stack to capture. We prioritized preserving temporal resolution of the videos and thus focused on the clearer planar view that captured cilia in profile. Raising ambient glucose from 1 to 11mM stimulated cilia movement by nearly a 2-fold increase in amplitude as measured by standard deviation of angle of deflection (θ), representing the transverse variance from the cilium base to a reference central axis (**Figure 1D, Movie S1**). Motility is most dynamic in the distal tip of the beta cell cilium, a region that is continually remodeled (*33*) and represents a specialized compartment enriched in GPCRs and other signaling molecules (*19, 27, 34*). High glucose induced greater axonemal bend and larger tip displacement, up to 10 µm, which exceeds that expected from purely thermal bending due to Brownian motion or actin-mediated internal forces (*13, 35*). Cilia beating frequency was higher in low glucose conditions as the cilia traversed shorter distances. All cilia waveform parameters, including curvature, frequency, periodicity, and amplitude, displayed increased distribution range under high glucose conditions, reflecting heterogeneity in beta cell responses. These observations suggest that active cilia motility may have evolved as a nutrient-seeking behavior to augment the detection of glucose.

We hypothesized that the mode of motility in beta cell cilia is conserved from that of conventional motile cilia, which requires specialized protein complexes organized throughout the axoneme. Therefore we examined human islets for endogenous expression of cilia motors and accessory components. Immunofluorescence revealed strong expression and ciliary localization of multiple motor proteins, including the spermatogenesis and flagellar assembly protein SPEF2, which is required for central pair formation (*36, 37*), the axonemal dynein intermediate-chain DNAI1, which forms the outer dynein arm and regulates motor activity (*38, 39*), the central pair-associated kinesin motor protein KIF9 (*40, 41*), and growth-arrest specific protein GAS8, which forms the nexin-dynein regulatory complex (N-DRC) (*42, 43*), all of which are essential for motile cilia and flagellar assembly and function (**Figure 2**). In human islets, DNAI1 and GAS8 are localized centrally and enriched in the proximal two-thirds of islet cilia, while SPEF2 and KIF9 are concentrated toward the distal tip, though all four proteins have detectable expression along the whole length of cilia.

**Figure 2.**
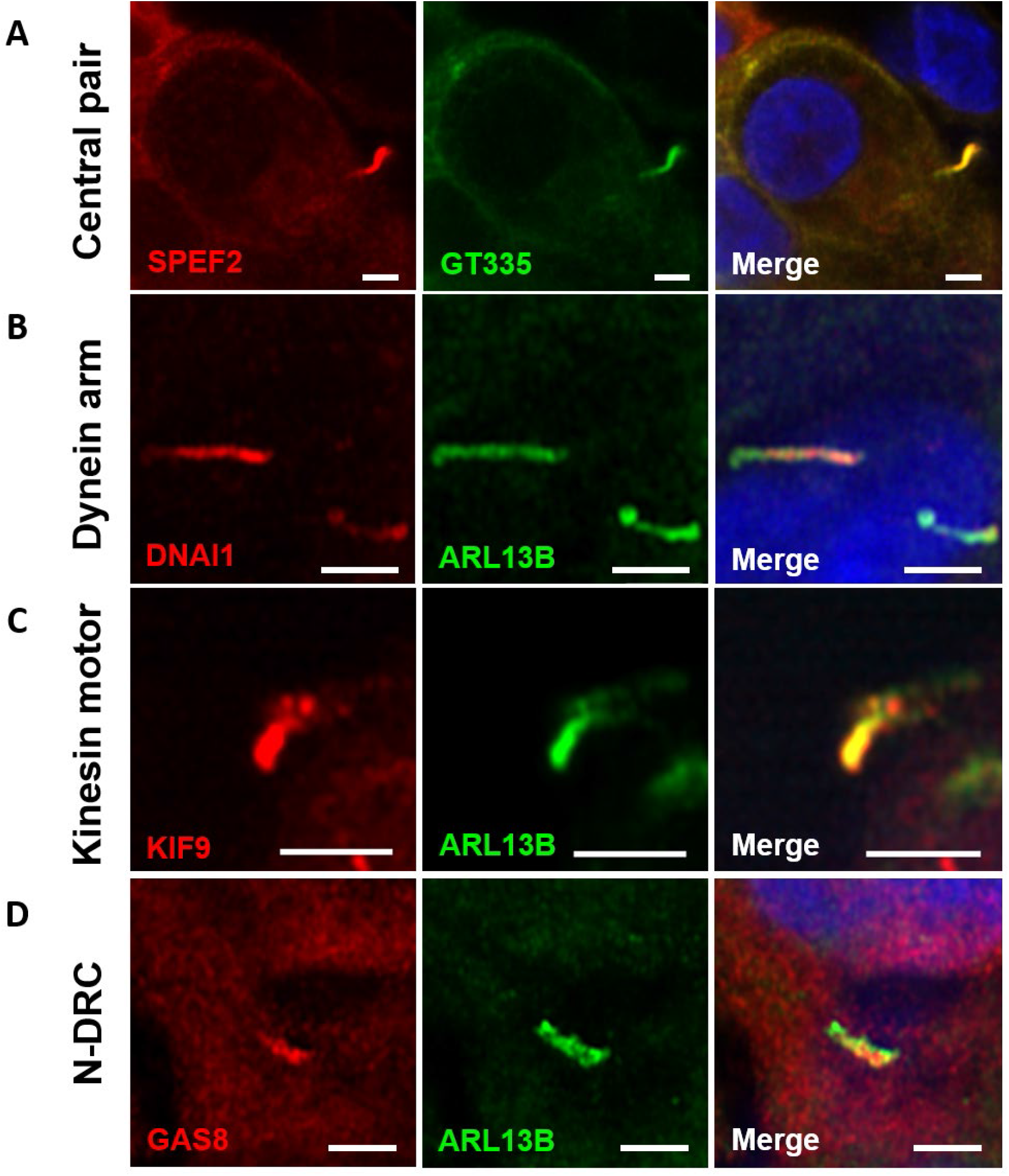
Endogenous motile cilia proteins in human islet cilia. (**A**) Healthy non-diabetic human islets were examined by confocal fluorescence microscopy using antibodies to SPEF2 (central pair-associated protein, red) and cilia marker GT335 (polyglutamylated tubulin, green). Co-localization of SPEF2 and cilia appears yellow in merged images. Nuclei are labeled with DAPI in blue. (**B-D**) The outer dynein arm protein DNAI1 (**B**, red), central pair protein KIF9 (**C**, red), and nexin-dynein regulatory complex (N-DRC) unit GAS8 (**D**, red) co-localize with cilia marker ARL13B (green). All scale bars, 2 µm.

To test the role of specific motor proteins in beta cell cilia motility, we used an array of pharmacologic inhibitors to block their function. Ciliobrevin D is a small molecule inhibitor of dynein ATPase (*44, 45*) and at 50 µM effectively inhibits cilia movement after 60 minutes of treatment (**Figure 3A, Movie S2**). Similar effects were seen using another dynein inhibitor, erytho-9-(2-hydroxynonyl) adenine (EHNA (*46*)), at 0.8 mM (**Figure 3B, Movie S3**). To confirm an energy requirement for ciliary motion, we performed ATP depletion using antimycin A and 2-deoxy-D-glucose, which profoundly inhibited cilia movement (**Figure 3C, Movie S4**). In contrast, inhibition of the actin-associated motor myosin II by blebbistatin did not obviously impair cilia movement (**Figure 3D, Movie S5**). These results suggest that cilia axonemal movement in beta cells depends on a microtubule-dynein driven mechanism and requires ATP hydrolysis.

**Figure 3.**
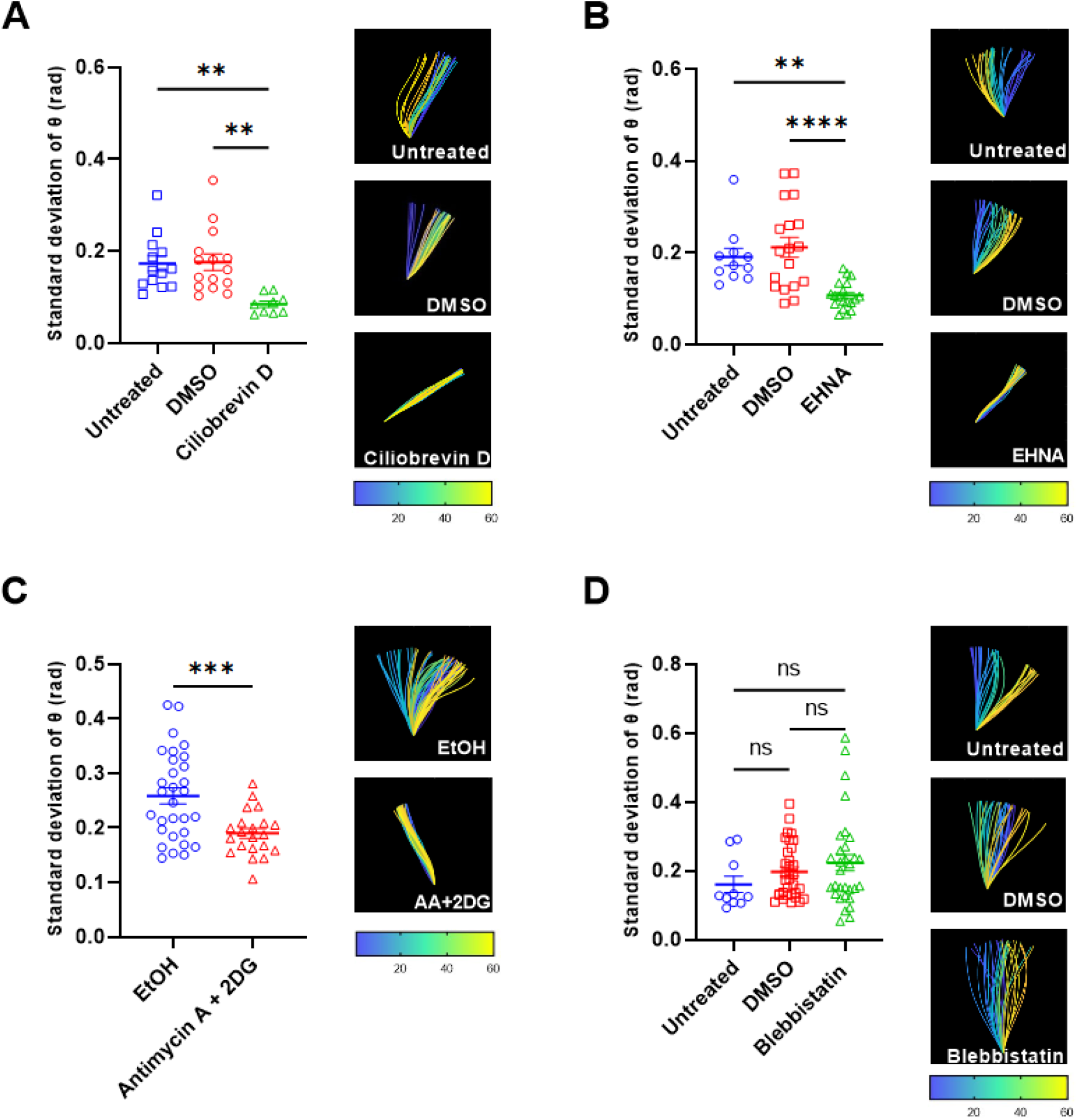
Beta cell cilia motility requires dynein, ATP, and glucose. (**A**) Cilia movement is inhibited by the dynein inhibitor ciliobrevin D, as shown in mean waveform amplitude (standard deviation of cilium tangent angle, θ, rads). DMSO-treated islet cilia exhibited similar motility as untreated islet cilia as an additional control. Representative wave traces are shown for each condition, temporal heatmap denoting the starting and ending frames of ciliary wave traces. (**B**) EHNA effectively inhibits cilia movement whereas the DMSO and untreated groups showed unaffected motility. (**C**) Antimycin A depletion of ATP and 2-deoxy-D-glucose (2DG) treatment inhibit cilia movement. (**D**) The actin-myosin inhibitor blebbistatin does not significantly affect ciliary beat amplitude. * \**p* < 0.01, * * \**p* < 0.001, * * * \**p* < 0.0001; n = 10-35 traces per condition, pooled from three or more independent experiments. Corresponding **Movies S2-5**.

Building on these findings, we next tested the relevance of dynein-mediated cilia motility to beta cell function. Dynein loss-of-function studies in intact human islets were performed using both pharmacologic and molecular approaches. As calcium is a crucial messenger underlying glucose-stimulated insulin secretion (GSIS), we first tested the effect of dynein inhibition on intracellular calcium dynamics. Glucose induces a large rise in cytosolic Ca^2+^ in normal beta cells, which is measurable by fluorescent sensors such as Calbryte 590 AM (**Figure 4A, Movie S6**). Treatment of whole islets with ciliobrevin D reduced both the time-onset and amplitude of the calcium influx (**Figure 4B, Movie S6**). This was corroborated by the effect of targeted knockdown of axonemal dynein DNAI1 on insulin secretion. Lentiviral knockdown in human islets resulted in over 80% decrease in *DNAI1* gene expression and nearly 50% decrease in GSIS (**Figure 4C**). Similarly, treatment of whole islets with ciliobrevin D completely arrested cilia motility and reduced insulin secretion by more than 50% in human islets (**Figure 4D**, average fold induction of 19.2 in DMSO vs. 9.3 in ciliobrevin D). Human islets also have motile cilia (**Movies S7-8**) and exhibited a range of insulin secretion in both baseline GSIS and response to ciliobrevin D, which was expected given donor heterogeneity. Together these results demonstrate that cilia motility is conserved between mouse and human islets and that insulin secretory function is at least in part dependent on cilia motility. Lastly, to test the hypothesis that islet primary cilia motility may be linked to human disease, we examined ciliary genes in individuals with type 2 diabetes. Using a previously published single-cell RNA-seq analysis of pancreatic islets obtained from human donors who were either healthy or diagnosed with type 2 diabetes (*47*), we manually curated a list of genes implicated in ciliary function from prior literature. When comparing average expression in alpha and beta cells separated by donor disease status, we observed that many cilia motility genes are more highly expressed in beta cells than other cell types, and higher in diabetic donors than healthy controls, including axonemal dynein and central pair assembly factors DNAH5, DNAAF2, SPAG16, and others (**Figure 4E**).

**Figure 4.**
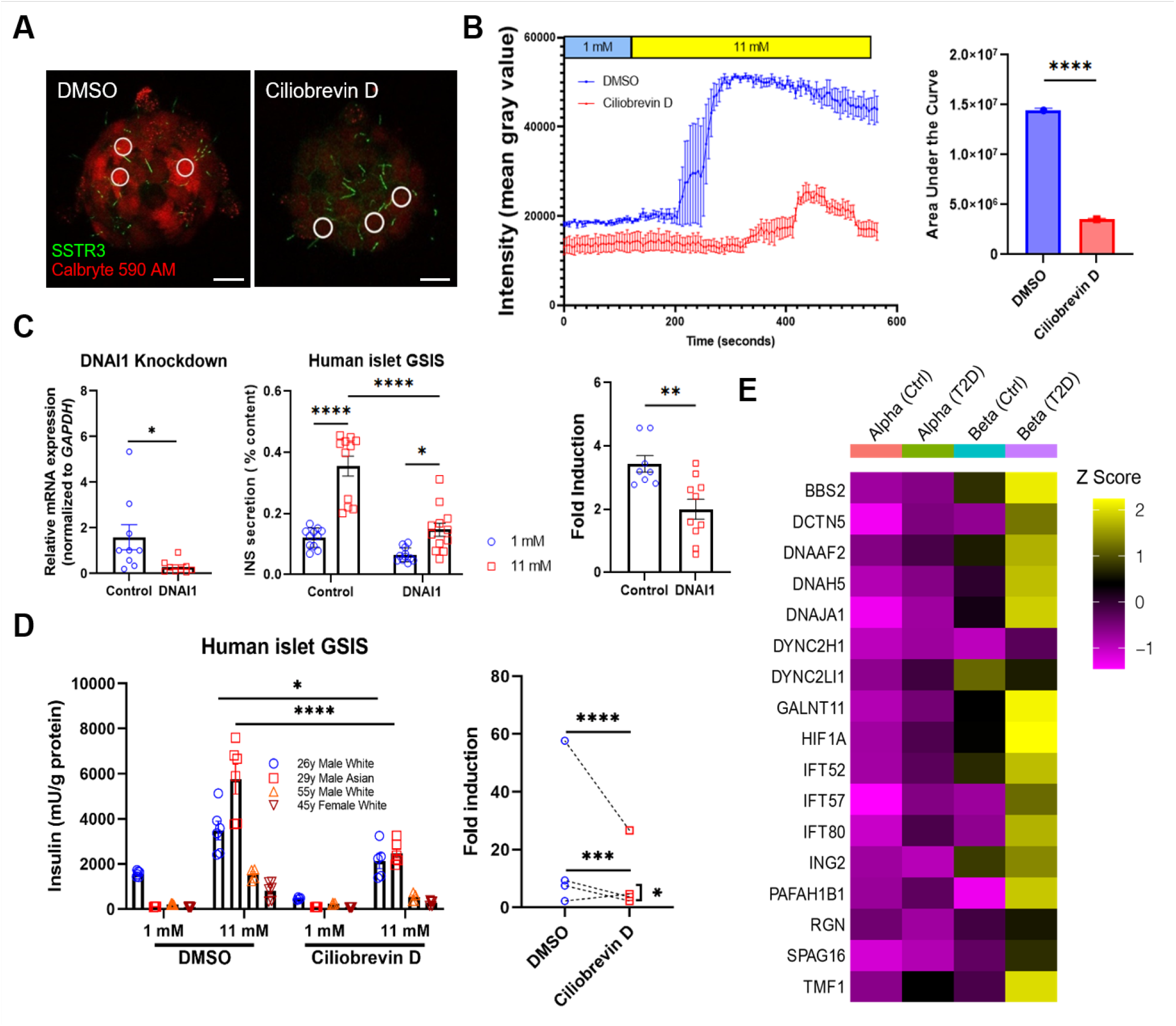
Cilia motility is required for glucose-stimulated insulin secretion. (**A**) Dual-color live imaging showing GFP-labeled beta cell cilia in whole islets labeled with red Calbryte dye (**Movie S6**). Still images show control vs. ciliobrevin-treated islets at peak calcium activity after glucose concentration was raised to 11 mM. Three representative regions of interest (ROIs) per islet are shown. Calbryte, red; cilia, green; scale, 20 µm. (**B**) Calcium response to glucose is delayed and ablated by ciliobrevin D, as shown by fluorescence intensity tracings and areas under the curve (AUC). Traces are composite of 3 ROIs and representative of islets examined in three independent experiments. (**C**) Dynein knockdown in human islets results in reduced *DNAI1* gene expression and diminished glucose-stimulated insulin secretion (GSIS); data plotted from two independent knockdown experiments. Statistical significance was analyzed by student t-test and two-way ANOVA. (**D**) Ciliobrevin D treatment inhibits GSIS in human islets. Fold induction by glucose was significantly reduced by ciliobrevin D; n = 4 donors. Statistical significance was analyzed by two-way ANOVA, *\*p < 0.05, * *p < 0.01, * * *p < 0.001, * * * *p< 0.0001*. (**E**) Cilia motility genes are more highly expressed in human T2D beta cells. Z-transformed normalized average expression of selected ciliary genes in alpha and beta cells from healthy controls and patients with T2D. Source data from a previously published single-cell RNA-seq study (*47*).

## DISCUSSION

The primary cilium is often called the cellular antenna for its prominent role in sensory perception and signaling. Here we show that the antenna is capable of active movement and that this facilitates islet beta cell nutrient sensing and hormone secretion. Motility of beta cell primary cilia is stimulated by glucose, the main metabolic signal for pancreatic islets. We find that islet primary cilia contain axonemal dynein motors and associated regulatory protein complexes, and functional studies confirm that their movement is driven by dynein via ATP hydrolysis, the classic force-generating system that confers cilia and flagellar motility. As dictated by islet topology, cells project their cilia either outward into the extra-islet space or inward to the interstitial space between adjacent cells (*16, 19, 48*). Therefore, cilia movement allows a 3D sampling of the exocrine-endocrine interface and of the islet interstitium, an important region where paracrine hormones, nutrient cues, and signaling molecules are present in high concentrations. Consistent with this idea, the mode of primary cilia motility appears to be a combination of the planar beating exhibited by conventional motile cilia and rotational movement of nodal cilia, such that each islet cell maximizes exploration of its surroundings and the collection of sensory stimuli. Similar non-periodic, 3D, wavelike motion provides exploratory function in other biological contexts, for example the hunting behavior of the unicellular predator *Lacrymaria olor* (*49*). Characterization of the complete 3D waveform (*50*) in beta cell primary cilia is therefore a priority of future research.

Ours is the first demonstration of dynein-controlled motility and the presence of motor accessory components in primary cilia. Primary cilia may move in the context of mechanosensory and signaling functions (*11, 13, 51*), but our discovery of motile primary cilia challenges the conventional classification of primary versus motile cilia. Our findings support a more permissive view where primary cilia might be classified as sensory-dominant but also motile. Accordingly, secondary cilia have been shown as motile-dominant while also capable of sensory and signaling function (*52*–*54*). There has been mounting evidence of primary cilia structural heterogeneity, where axonemal architecture can deviate from the classic “9+0” arrangement (*17, 18, 55*). Our current study shows the presence of dynein arms and central pairs in beta cell primary cilia. With the advance of electron tomography and improved sample preservation techniques, we expect to soon see primary cilia ultrastructure that demonstrates microtubule central pairs and dynein regulatory complexes, labile structures whose detection is critically dependent on fixation protocol (*55, 56*).

Based on the cilia motility behavior we observed in live islets and the strong link between cilia movement and insulin secretion, we propose that cilia motility is an essential component of beta cell function. This notion is consistent with the observation that cilia motility genes are increased in the beta cells of people with type 2 diabetes, a disorder often characterized by insulin hypersecretion. In islets, where there is heterogeneity among cilia orientation and alignment, having motile cilia is likely important for intercellular communication as neighboring cilia may physically interact to transduce signals between cells. The specific configuration and movement of individual cilia is therefore of interest, and future studies might delineate a functional role of cilia motility in mediating paracrine or juxtacrine signaling. Whether motility in beta cell cilia induces mechanotransduction and Ca^2+^ signaling through TRPV and polycystin channels (*57*) is another open question. The demonstration of motility genes across human islet transcriptome studies suggests that primary cilia motility is conserved across different cell types and likely mediates islet-specific function. The finding of cilia motility gene changes in human T2D also suggests a role for beta cell cilia motility in diabetes pathophysiology and may provide a new therapeutic target to modulate insulin secretion.

## Supporting information

Movie S1. Beta cell cilia movement in low vs high glucose

Movie S2. Ciliobrevin inhibits beta cell motility

Movie S3. EHNA inhibits beta cell cilia motility

Movie S4. AA2DG inhibits beta cell cilia motility

Movie S5. Blebbistatin does not affect beta cell cilia motility

Movie S6. Ciliobrevin disrupts calcium dynamics in cilia reporter beta cells

Movie S7. Human islets with green cADDis sensors

Movie S8. Human islet cilia with siR-Tubulin

## SUPPLEMENTARY MATERIALS

### MATERIALS AND METHODS

#### Generation of beta-cell cilia GFP mice

INS1-Cre mice from Jackson Laboratories (JAX #026801) were crossed with SSTR3 GFP mice (*29*) kindly provided by Yoder Lab at UAB. Mice at weaning were genotyped by a commercial vendor (Transnetyx) and fed a standard rodent diet (PicoLab Mouse Diet S053, 13.2% calories from fat). Both male and female mice were used for experiments between 2 and 4 months of age. Animals were maintained in accordance with Institutional Animal Care and Use Committee regulations at the Washington University School of Medicine, approval #20190188.

#### Human islets

Intact islets from cadaveric non-diabetic human donors were obtained from the NIDDK Integrated Islet Distribution Program (IIDP) or purchased commercially from Prodo Laboratories, Inc. Upon arrival, islets were visually inspected, washed twice in islet media (RPMI-1640, 10% FBS, 11 mM glucose, 1% (v/v) penicillin/streptomycin, and 20 mM HEPES), and cultured overnight prior to experiments. Islets from both male and female donors were used. All references to “low” and “high” glucose in insulin secretion and imaging experiments correspond to 1 and 11 mM glucose, respectively.

#### Electron microscopy

Human islet cilia were examined using electron microscopy. Isolated islets were fixed in Karnovsky’s solution (3% glutaraldehyde, 1% paraformaldehyde), then secondarily fixed in osmium tetroxide, dehydrated in alcohol, embedded in resin, and polymerized at 90 °C for 48 hours. Ultrathin sections of 90 nm thickness were cut and stained with uranyl acetate and imaged with a JEOL 1200 EX.

#### Live cilia motility imaging

Islets were isolated from beta-cell cilia GFP mice using an established protocol (*58*) and recovered for 24 hours in islet media at 37 °C and 5% CO_2_. On imaging day, islets were treated for 1 hour with inhibitors and vehicle control, including ciliobrevin D (50 µM in DMSO, Sigma 250401), EHNA (0.8 mM in DMSO, Cayman Chemical 13352), antimycin A (20 µM in EtOH, Sigma A8674), 2-deoxy-D-glucose (10 mM in PBS, Sigma D8375), and blebbistatin (50 µM in DMSO, Sigma 203389). Inhibitor treatments were performed at 1 mM glucose except antimycin A and 2-deoxy-D-glucose studies, which had no glucose supplementation. Treatment conditions were well-tolerated by cells as determined by viability assays. For human islet cilia live imaging, siR-Tubulin (Cytoskeleton CY-SC002) and green cilia-targeted cADDis sensors (Montana Molecular D0201G) were used to label cilia in intact human islets. After treatment or labeling, islets were washed twice with KRBH and transferred to 35 mm glass-bottom imaging dishes. Live cilia motility images were recorded using an inverted Zeiss LSM880 fluorescence microscope (Nikon, Ti-E) using a 63x oil immersion objective. Each experiment was independently performed by 2-4 lab members to ensure reproducible findings. Imaging parameters varied among experiments and researchers but generally included 60-120 cycles, 7-12 slices (for z-stack images only), 1-2 AU (pinhole size), 512×512 pixel size, 16 bit depth, 0.26-0.44 µm/pixel, and up to 660 seconds duration.

#### Cilia waveform analysis

Cilium traces were obtained using a custom program written in MATLAB (The MathWorks Inc., Natick, MA). Using the first image of a time series, the user manually traces the cilium to obtain length, base coordinates, and base angle; the program then automatically traces the path of the cilium in successive frames, using the following algorithm. The length of the cilium is divided into *n* segments. For the first segment, a rectangular array of points spanning the cilium width and the length of one segment is rotated through a range of angles about the cilium base. At each angle, the weighted average of the interpolated pixel intensities at all points in the array is calculated. The next point along the cilium is located using this angle, the base coordinate, and the segment length. To improve the robustness of the angle selection, terms are applied to penalize curvature, translational velocity, rotational velocity, and change in curvature per time step. The process is then repeated along successive points along the cilium. When all *n*+1 points along the cilium have been identified, the program proceeds to the next image in the sequence.

The program requires that any translation of the base of the cilium be removed beforehand (*30*). Quantities of interest including period, periodicity, curvature, and amplitude are calculated by post-processing the raw angle data obtained above. Oscillation period is estimated from the nonzero time delay that maximizes the autocorrelation of the angle data; this delay is found at all points along the cilium and averaged to obtain a single estimate of the period. The mean of the maximum autocorrelation value at each point is used as a measure of periodicity, or how similar the beats are to each other. For each frame, a fourth order polynomial is fit to the angle data and used to reconstruct angle as a continuous function of the arc length, *s*. The shape of the cilium, as represented by *x* and *y* positions along the cilium axis, is reconstructed by integrating the cosine and sine, respectively, of the angle along the length of the cilium. Curvature is calculated as the derivative of the fitted angle function with respect to arclength. Beating amplitude is quantified by taking the standard deviation of all angle data within the middle 80% of the length of the cilium, excluding the base and the tip.

#### Calcium imaging in intact mouse islets

Intracellular calcium was measured using the cell-permeant dye Calbryte 590 AM (AAT Bioquest, CA). Islets were incubated in 4 µM Calbryte 590 AM at 37 °C. Mouse and human islets were loaded for 15 and 30 minutes, respectively. Following dye incubation, islets were washed and allowed to recover in KRBH buffer for 10 minutes. Islets were then transferred to a 4-chamber glass-bottom imaging dish and mounted on a climate-controlled stage with 37 °C and 5% CO_2_. Calcium imaging was performed on a Zeiss LSM880 inverted confocal system using a 63x oil immersion objective. Calbryte 590 AM was excited by a 561 nm laser and detected in the range of 571 - 700 nm. Baseline recordings were performed at 1 mM glucose for 5 minutes, then glucose was added to a final concentration of 11 mM by manual pipetting, and islets were continuously imaged for a total duration of 12 minutes. 16-bit 512×512 pixel images were acquired every 465 ms, which is the frequency best suited to resolve islet Ca^2+^ dynamics upon glucose stimulation. For two-channel z-stacked time-lapse imaging of cilia overlapped with calcium signals, 12 µm-thick islet regions were imaged in 7 consecutive planes along the z-axis with a total scan time of 3.25 seconds per cycle. A total of 120 cycles of z-stack time-lapse were recorded and glucose was added on cycle 30 (140 seconds) while islets were continuously imaged for a total duration of 564 seconds. Calcium traces were extracted using Calbryte fluorescence integrated intensity (mean gray value) from ROIs in ImageJ software (FIJI, (*59*)). Traces were selected from cells that exhibited minimal motion artifacts and were averaged across biological replicates. Calcium kinetics were time-corrected in Excel and then plotted in Prism Graphpad software (San Diego, CA).

#### Immunohistochemistry

Isolated islets were washed with PBS and fixed with 4% paraformaldehyde (PFA) for 30 minutes and permeabilized with 0.3% Triton X-100 in PBS (PBST) for 30 minutes at room temperature. After incubation with blocking buffer PBS with 10% normal goat serum for 1 hour at room temperature, islets were incubated for 48 hours at 4 °C with primary antibodies diluted in PBST. The next day, islets were washed, incubated overnight at 4 °C with secondary antibodies diluted in PBS, and washed again with PBS. DAPI provided nuclear counterstain. Islets were mounted on glass slides with ProLong Gold Antifade Mountant (Thermo Fisher P36930) prior to imaging. Primary antibodies used for cilia immunostaining include: ARL13B (Proteintech 17711-1-AP, NeuroMab 75-287), AcTUB (Proteintech 66200-1-Ig, Sigma-Aldrich T7451), GFP (Thermo Fisher Invitrogen #A-11122), Polyglutamylated Tubulin (GT335 AdipoGen AG-20B-0020), DNAI1 (NeuroMab 75-372), GAS8 (Atlas Antibodies HPA041311), SPEF2 (Atlas Antibodies HPA040343), and KIF9 (Atlas Antibodies HPA022033). Secondary antibodies include Alexa Fluor goat-anti-mouse and goat-anti-rabbit (Invitrogen).

Images were acquired on a Zeiss LSM 880 microscopy with AiryScan detector (Zeiss, Oberkochen, Germany). Plan-Apochromat 63x /1.40NA Oil DIC M27 oil immersion objective was used for all experiments. Images were acquired using an optical magnification of at least 1.8x. XY pixel size was optimized per Zeiss Zen software (11 nm pixel size). Z pixel dimension was set to 250 nm per Z-step. Laser power and Z-ramp (when necessary) for each channel was set independently to equalize signals at the top and bottom of the imaging stack and to occupy the first ∼1/3 of the detector dynamic range. Scan speed was set to 6, scan averages to 2, gain to 700-900, digital offset to 0, and digital gain to 1. Confirmation of AiryScan detector alignment was performed before image acquisition and was rechecked with every new slide. Following acquisition, images were 3D Airyscan processed in Zeiss Zen Black.

#### Insulin secretion

For DNAI1 knockdown experiments, after successful lentiviral infection and incubation in islet media, ten size-matched, re-aggregated human islets were picked from 384-well microplates and equilibrated in the Krebs-Ringer Bicarbonate HEPES (KRBH) buffer (128 mM NaCl, 4.8 mM KCl, 1.2 mM KH_2_PO_4_, 1.2 mM MgSO_4_ 7H_2_O, 2.5 mM CaCl_2_, 20 mM HEPES, 5 mM NaHCO_3_, and 0.1% BSA, pH 7.4) at 2.8 mM glucose for 1 hour at 37 °C. Islets were then transferred to the KRBH buffer with 1 or 11 mM glucose respectively for 1 hour at 37 °C for insulin secretion. Supernatant and islets were collected and stored at -80 °C before quantification using Lumit Insulin Immunoassay Kit (Promega 2021). Secreted insulin levels were normalized to total insulin content of whole islets after acid ethanol extraction. Fold induction was calculated as the ratio of insulin secretion at 11 mM glucose vs. 1 mM glucose.

For ciliobrevin experiments, five size-matched human islets were equilibrated in triplicate in 24-well plates containing 400 µL KRBH buffer at 1 mM glucose for 1 hour at 37 °C. Islets were then transferred to 1 or 11 mM glucose for secretion for 1 hour at 37 °C, with or without ciliobrevin D (50 µM, DMSO vehicle control). Supernatants and islets were collected separately and stored at -80 °C until insulin measurements, performed on Crystal Chem insulin ELISA assay kits according to manufacturer specifications. Secreted insulin was normalized to total protein, which was determined using a Pierce BCA protein assay kit (Thermo Scientific 23225) according to the manufacturer’s instructions. Fold induction was calculated as the ratio of insulin secretion at 11 mM glucose vs. 1 mM glucose.

#### Generation of lentiviral knockdown human islets

Human lentiviral particles were generated from Origene (TL313431V). After dissociation of human islets using Accutase (Innovative Cell Technologies AT-104) for 20 minutes at 37 °C, cells were seeded in microwells or low-cell attachment plates and treated with control and DNAI1 shRNA GFP lentiviral particles at the concentration of 1×10^6^ titer unit/mL for 24-48 hours. These samples were further incubated for 96 hours to allow recovery and re-aggregation, then used for gene expression and insulin secretion experiments.

#### Quantitative real time PCR

After shRNA knockdown, re-aggregated human islets were lysed in 350 µL lysis buffer RA1 with β-mercaptoethanol per 100-200 islets. RNA was purified with a NucleoSpin RNA kit (Macherey-Nagel, Germany) and cDNA was synthesized using a Thermo Fisher High-Capacity cDNA reverse transcription kit at 200 ng/20 µL. qPCR was performed in triplicate on a 7900 Step One Plus RT-PCR machine (Applied Biosystems) using 2x PowerSYBR Green PCR Master Mix (Thermo Fisher 4367659). Changes in gene expression were quantified using 2^-Δ ΔCt^, and results were normalized to the housekeeping gene GAPDH. The following primer sequences were used: human *DNAI1*: F: 5’-TCAGCCAAGTCTGGCAAGCACT-3’, R: 5’- GAGTCCAAGACACAATCCTGCC-3’; human *GAPDH*: F: 5’- GTCTCCTCTGACTTCAACAGCG-3’, R: 5’-ACCACCCTGTTG CTGTAGCCAA-3’.

#### Bioinformatics

Raw read counts from the previously published single-cell RNA-seq data of human pancreatic islet cells was downloaded from ArrayExpress (Accession: E-MTAB-5061). All data analysis was performed in R using Seurat (version 4.0.5). Cells with at least 1,500 expressed genes and at least 15,000 detected unique molecular identifiers were retained for analysis. Counts were log-normalized and scaled, and clustering was performed by UMAP. Cell clusters were identified by marker gene expression. Genes involved with ciliary function were identified by literature review and a subset of these were identified with measurable expression and variability across cell types in the dataset.

#### Statistics

Data are presented as mean ± SEM. Statistical significance was analyzed by paired student t-test (2 groups) or ANOVA (>2 groups), *\*p < 0.05, * *p < 0.01, * * *p < 0.001, * * * *p< 0.0001*. Sample size and number of replicates for each experiment are indicated in figure legends.

## ACKNOWLEDGMENTS

We are grateful to the Yoder lab at UAB and the Piston, Brody, Horani, and Millman labs at Washington University for reagent sharing. We thank X. W. Ng for islet imaging expertise and pilot studies. We thank the Hughes laboratory members and the Washington University cilia research community for data review as well as S. Brody and C. Semenkovich for critical reading of the manuscript. All graphics were created with BioRender.com.

## Author contributions

Conceptualization: JWH, JHC, HFR

Methodology: JWH, JHC, ZAL, HFR, TU, LW, PVB

Investigation: JWH, JHC, ZAL, LZ, BDM, HFR, TU, LW

Visualization: JWH, JHC, ZAL, LZ, BDM

Funding acquisition: JWH, BDM, PVB

Supervision: JWH, PVB

Writing - original draft: JWH, JHC, ZAL

Writing - review & editing: All authors

## Data and materials availability

All data are available in the main text and supplementary materials. Transgenic animals will be made available under material transfer agreements. Code used to analyze the publicly available single-cell RNA-seq data is available upon request.

## SUPPLEMENTAL DATA

### Movie files Mouse islet imaging

- **S1. Beta cell cilia movement in low vs high glucose**
- **S2. Dynein inhibitor ciliobrevin blocks beta cell cilia motility**
- **S3. Dynein inhibitor EHNA blocks beta cell cilia motility**
- **S4. ATP depletion by AA2DG inhibits beta cell cilia motility**
- **S5. Actin inhibitor blebbistatin does not affect beta cell cilia motility**
- **S6. Ciliobrevin disrupts calcium dynamics in cilia reporter beta cells**

### Human islet imaging

- **S7. Human islet cilia motility as imaged using green cilia cADDis**
- **S8. Human islet cilia motility as imaged using siR-Tubulin**

## LEGENDS FOR MOVIE FILES

**Movie S1. Beta cell cilia movement in low vs. high glucose**.

Video images of beta cell cilia on intact murine islets incubated in 1 mM glucose were captured with a frame time of 465.45 ms. Glucose level was raised to 11 mM and cells were re-imaged, showing increased cilia motility as measured by amplitude. Scale bar = 5 µm.

**Movie S2. Dynein inhibitor ciliobrevin blocks beta cell cilia motility**.

Video images of beta cell cilia on mouse islets after 1-hour treatment with 0.5% DMSO or 50 µM ciliobrevin D in 1 mM glucose-containing KRBH were recorded with a frame time of 20.26 seconds, showing inhibition of beta cell cilia motility by ciliobrevin D. Scale bar = 5 µm.

**Movie S3. Dynein inhibitor EHNA blocks beta cell cilia motility**.

Video images of beta cell cilia on mouse islets incubated in DMSO or 80 mM EHNA in 1 mM glucose-containing KRBH for 1 hour were captured with a frame time of 20.26 seconds, showing that beta cell cilia motility was blocked by EHNA. Scale bar = 5 µm.

**Movie S4. ATP depletion by AA2DG blocks beta cell cilia motility**.

Video images of beta cell cilia on mouse islets incubated in EtOH or 20 µM Antimycin A and 10 mM 2-deoxy-D-glucose in PBS with no glucose-containing KRBH for 1 hour were recorded with a frame time of 1.27 seconds, demonstrating that beta cell cilia motility requires ATP and glucose. Scale bar = 5 µm.

**Movie S5. Actin inhibitor blebbistatin does not affect beta cell cilia motility**.

Video images of beta cell cilia on whole mouse islets incubated in DMSO or 50 µM blebbistatin in 1 mM glucose-containing KRBH for 1 hour were captured with a frame time of 20.26 seconds, showing that beta cell cilia motility does not depend on the actin motor myosin II. Scale bar = 5 µm.

**Movie S6. Ciliobrevin disrupts calcium dynamics in cilia reporter beta cells**.

Video images of beta cell cilia and calcium influx in INS1xSSTR3-GFP mouse islets labeled with 4 µM Calbryte 590 AM in islet media were captured using two-channel time-lapse and z-stack imaging with a frame time of 465.45 ms. Glucose concentration was raised from 1 mM to 11 mM during image capture, and calcium flashes indicate cell response to glucose. Video shows maximum intensity projection; some cilia appear segmented due to limited z-plane sampling. Scale bar = 50 µm.

**Movie S7. Human islet cilia motility as imaged using green cilia cADDis**

Video images of spontaneous cilia movements on human islets were labeled overnight with green cilia cADDis in islet media. Images were recorded at 1 mM glucose with a frame time of 2.53 seconds. Scale bar = 2 µm.

**Movie S8. Human islet cilia motility as imaged using siR-Tubulin**

Video images of human islet cilia labeled with 1 µM siR-Tubulin and 10 µM verapamil in islet media for 1 hour were captured with a frame time of 1.27 seconds, showing that human islet cilia move. Scale bar = 2 µm.

